# GimmeMotifs: an analysis framework for transcription factor motif analysis

**DOI:** 10.1101/474403

**Authors:** Niklas Bruse, Simon J. van Heeringen

**Affiliations:** Radboud University, Faculty of Science, Department of Molecular Developmental Biology, Radboud Institute for Molecular Life Sciences, 6500 HB Nijmegen, The Netherlands

## Abstract

**Background:** Transcription factors (TFs) bind to specific DNA sequences, TF motifs, in cis-regulatory sequences and control the expression of the diverse transcriptional programs encoded in the genome. The concerted action of TFs within the chromatin context enables precise temporal and spatial expression patterns. To understand how TFs control gene expression it is essential to model TF binding. TF motif information can help to interpret the exact role of individual regulatory elements, for instance to predict the functional impact of non-coding variants.

**Findings:** Here we present GimmeMotifs, a comprehensive computational framework for TF motif analysis. Compared to the previously published version, this release adds a whole range of new functionality and analysis methods. It now includes tools for *de novo* motif discovery, motif scanning and sequence analysis, motif clustering, calculation of performance metrics and visualization. Included with GimmeMotifs is a non-redundant database of clustered motifs. Compared to other motif databases, this collection of motifs shows competitive performance in discriminating bound from unbound sequences. Using our *de novo* motif discovery pipeline we find large differences in performance between *de novo* motif finders on ChIP-seq data. Using an ensemble method such as implemented in GimmeMotifs will generally result in improved motif identification compared to a single motif finder. Finally, we demonstrate *maelstrom*, a new ensemble method that enables comparative analysis of TF motifs between multiple high-throughput sequencing experiments, such as ChIP-seq or ATAC-seq. Using a collection of ~200 H3K27ac ChIP-seq data sets we identify TFs that play a role in hematopoietic differentiation and lineage commitment.

**Conclusion:** GimmeMotifs is a fully-featured and flexible framework for TF motif analysis. It contains both command-line tools as well as a Python API and is freely available at: https://github.com/vanheeringen-lab/gimmemotifs.

## Introduction

The regulatory networks that determine cell and tissue identity are robust, yet remarkably flexible. Transcription factors (TFs) control the expression of genes by binding to their cognate DNA sequences, TF motifs, in cis-regulatory elements [1]. To understand how genetic variation affects binding and to elucidate the role of TFs in regulatory networks we need to be able to accurately model binding of TFs to the DNA sequence.

The specificity of DNA-binding proteins can be modeled using various representations [2]. One of the most widely adopted is the position frequency matrix (PFM). This matrix, a TF motif, contains (normalized) frequencies of each nucleotide at each position in a collection of aligned binding sites. These PFMs can be derived from high-throughput experiments such as Chromatin Immunoprecipitation followed by sequencing (ChIP-seq) [3,4,5,6], HT-SELEX [7] or Protein Binding Microarrays (PBMs) [8]. Through straightforward transformations a PFM can be expressed as a weight matrix, using log likelihoods, or information content, using the Kullback-Leibler divergence.

We previously published GimmeMotifs, a *de novo* ChIP-seq motif discovery pipeline [9]. Here, we present a new and updated version of the GimmeMotifs software. Compared to the previous version, it now contains a whole range of Python modules and command-line tools to provide an comprehensive framework for transcription factor motif analysis. Amongst other possibilities it can be used to perfom *de novo* motif analysis, cluster and visualize motifs and to calculate enrichment statistics. A new ensemble method, *maelstrom*, can be used to determine differential motif activity between multiple different conditions, such as cell types or treatments. We illustrate the functionality of GimmeMotifs using three different examples.

## Findings

GimmeMotifs includes several different modules (Table 1), that can all be used either via a command-line tool or using the Python API. In the next sections we illustrate the functionality using three different examples.

**Table 1:**
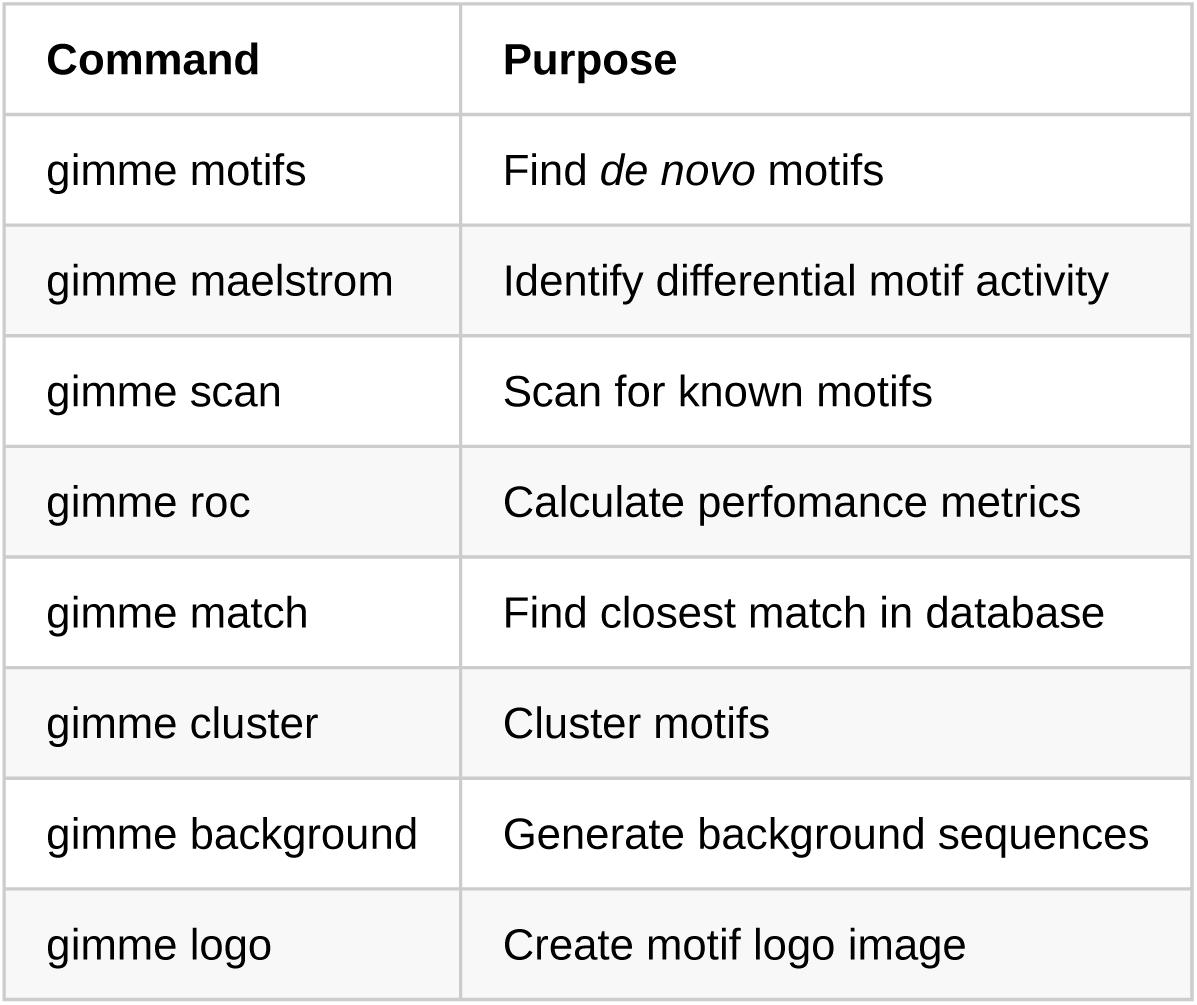
Command line tools included with GimmeMotifs. All these tools can also be used via the GimmeMotifs API.

| Command | Purpose |
| --- | --- |
| `gimme motifs` | Find *de novo* motifs |
| `gimme maelstrom` | Identify differential motif activity |
| `gimme scan` | Scan for known motifs |
| `gimme roc` | Calculate performance metrics |
| `gimme match` | Find closest match in database |
| `gimme cluster` | Cluster motifs |
| `gimme background` | Generate background sequences |
| `gimme logo` | Create motif logo image |

### Benchmark of transcription factor motif databases

A variety of transcription factor (TF) motif databases have been published based on different data sources. One of the most established is JASPAR, which consists of a collection of non-redundant, curated binding profiles [10]. The JASPAR website contains many other tools and the underlying databases are also accessible via an API [11]. Other databases are based on protein binding micorarrays [8], HT-SELEX [7] or ChIP-seq profiles [3,4,5,6]. CIS-BP integrates many individual motif databases, and includes assignments of TFs to motifs bases based on DNA binding domain homology [12].

For the purpose of motif analysis, it is beneficial to have a database that is non-redundant (i.e., similar motifs are grouped together), yet as complete as possible (i.e., covers a wide variety of TFs). To establish a quantitative measure of database quality, we evaluated how well motifs from different databases can classify ChIP-seq peaks from background sequences. In this benchmark, we used randomly selected genomic sequence as background. The following databases were included in our comparison: JASPAR 2018 vertebrate [10], SwissRegulon [13], Homer [6], Factorbook [3], the ENCODE motifs from Kheradpour et al. [4], HOCOMOCO [5], the RSAT clustered motifs [14] and the IMAGE motif database created by Madsen et al. [15]. We compared these databases to the non-redundant vertebrate motif database included with GimmeMotifs (v5.0, see Methods).

As a reference data set we downloaded all ChIP-seq peaks from ReMap 2018 [16], and selected the TFs with at least 1,000 peaks. When there were more than 5,000 peaks for a TF we randomly selected 5,000 peaks for the analysis.

We then evaluated the eight motif databases to test how well they could discriminate TF peaks from random genomic sequences using the GimmeMotifs tool `gimme roc`. This tool calculates a range of performance metrics to compare motif quality. For each TF and motif database combination we selected the single best performing motif, depending on the metric under consideration.

Figure 1A shows distribution of the ROC AUC (area under the curve for the Receiver Operator Curve) of the best motif per database for all 294 transcription factors in a box plot. The ROC curve plots the fraction of true positives (TPR, or sensitivity) against the fraction of false positives (FPR, or 1 - specificity). The ROC AUC value will generally range from 0.5 (no improvement on random guessing) to 1.0 (perfect classifier). As Figure 1A illustrates, the collection of TFs generally shows a wide distribution of ROC AUCs. For some factors, such as ELK1, CTCF, CBFB and MYOD1, peaks are relatively easy to classify using a single PFM motif. Other factors do not have peaks with a consistently enriched motif, or do not contain a sequence-specific DNA-binding domain, such as EP300 or CDK2 for example.

**Figure 1:**
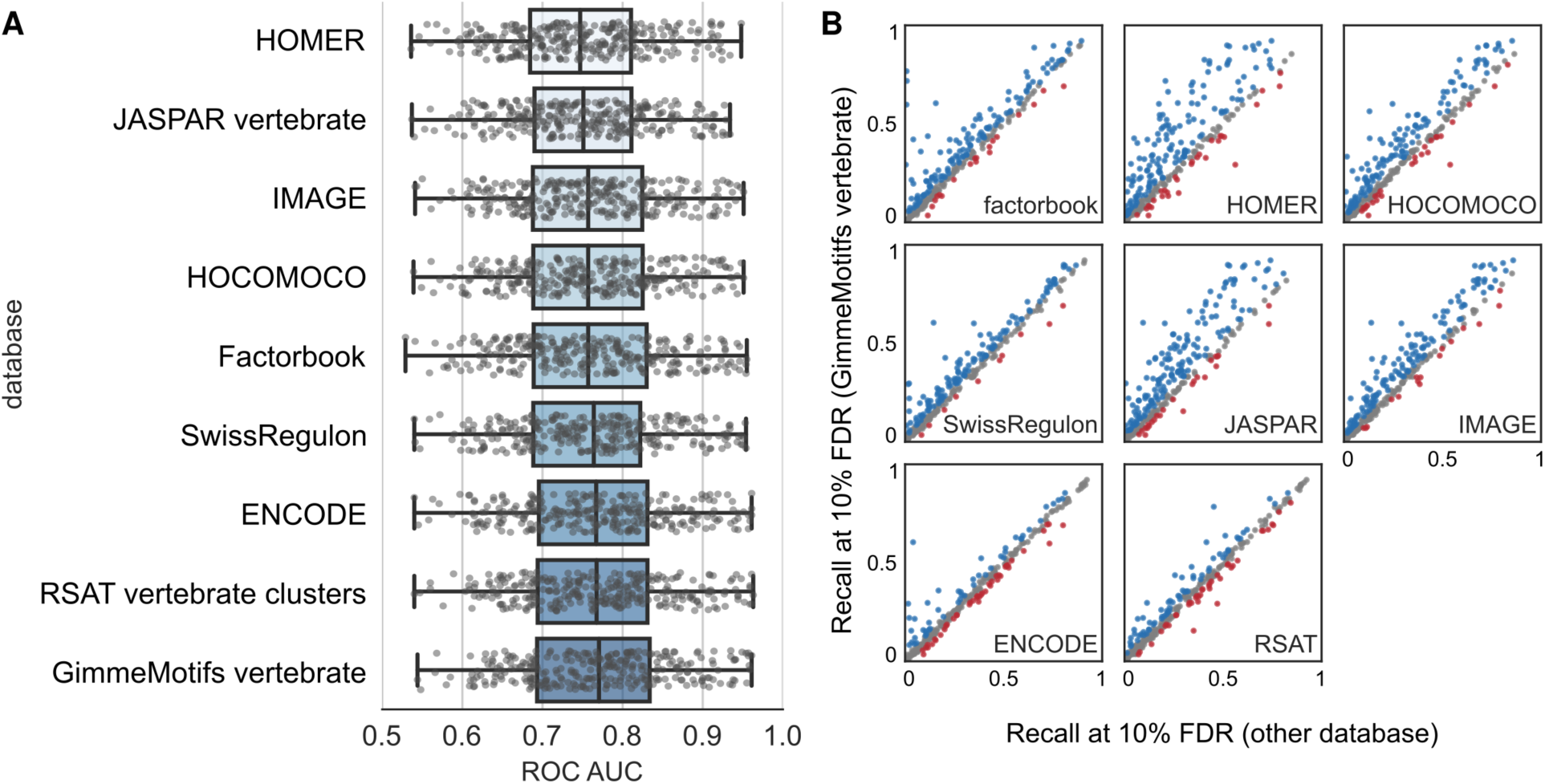
Benchmark of transcription factor motif databases. **A)** Motif-based classification of binding sites for 294 TFs from the ReMap ChIP-seq database. For all TFs 5,000 peaks were compared to background regions using each motif database. The boxplot shows the the ROC AUC of the best motif per database for all TFs. Every point in this plot is based on one TF ChIP-seq peak set. **B)** Recall at 10% FDR of motif databases compared to the GimmeMotifs vertebrate motif database (v5.0). The same data is used as in **A)**. The X-axis represents the recall for the different databases, the Y-axis represents the recall for the GimmeMotifs vertebrate database. Differences of more than 0.025 are marked blue, and less then −0.025 red.

The difference in maximum ROC AUC between databases is on average not very large, with a mean maximum difference of 0.05. The largest difference (~0.24) is found for factors that were not assayed by ENCODE, such as ONECUT1, SIX2 and TP73, and are therefore not present in the Factorbook motif database. Unsurprisingly, the databases that were based on motif collections of different sources (ENCODE, IMAGE, RSAT and GimmeMotifs) generally perform best. It should be noted that, for this task, using motif databases based on motif identification from ChIP-seq peaks is in some sense “overfitting”, as the motifs in these databases were inferred from highly similar data.

While the ROC AUC is often used to compare the trade-off between sensitivity versus specificity, in this context it is not the best metric from a biological point of view. An alternative way of measuring performance is evaluating the recall (i.e. how many true peaks do we recover) at a specific false discovery rate. This is one of the criteria that has been used by the ENCODE DREAM challenge for evaluation [17]. Figure 1B shows scatterplots for the recall at 10% FDR for all motif databases compared to the clustered, non-redundant databases that is included with GimmeMotifs. This database shows better performance than most other databases using this benchmark. The non-redundant RSAT database, which was created in a very similar manner [14], scores comparably.

These results illustrate how `gimme roc` can be used for evaluation of motifs. The choice of a motif database can greatly influence the results of an analysis. The default database included with GimmeMotifs shows good performance on the metric evaluated here. However, this analysis illustrates only one specific use case of application of a motif database. In other cases well-curated databases such as JASPAR can be beneficial, for instance when linking motifs to binding proteins. Any of these databases can be easily used in all GimmeMotifs tools.

### Large-scale benchmark of *de novo* motif finder performance on ChIP-seq peaks

It has been noted that there is no *de novo* motif prediction algorithm that consistently performs well across different data sets [18]. New approaches and algorithms for *de novo* motif discovery continue to be published, however, many of them are not tested on more than a few datasets. Benchmarks that have been published since Tompa et al. [19,20] typically have tested only a few motif finders or used only a few datasets.

Here, we used the GimmeMotifs framework as implemented in `gimme motifs` to benchmark 14 different *de novo* motif finders. To evaluate the different approaches, we downloaded 495 peak files for 270 proteins from ENCODE [21] and selected the 100bp sequence centered on the summit of top 5,000 peaks. These will be the peaks most likely to contain the primary TF motif, and should provide a straightforward test-case for the *de novo* motif finders. Ranking and selecting peaks in this manner is a widely adopted practice and we use this procedure also for our benchmark.

However, when analyzing ChIP-seq data in detail, it might be preferable to analyze to full complement of peaks.

Of the top peaks, half were randomly selected as a prediction set and the other half was used for evaluation. As a background set we selected regions of the same length flanking the original peaks. This will account for sequence bias according to genomic distribution. To assess the performance, we calculated two metrics, the ROC AUC and the recall at 10% FDR. Figure 2A shows the distribution of the ROC AUC scores over all ENCODE peaks in a boxplot, ordered by the mean ROC AUC. The ROC AUC is distributed between 0.58 and 0.98, with a mean of 0.75. All proteins that have low ROC AUC are not sequence-specific transcription factors such as POL2, TAF7 and GTF2B, the PRC2-subunit SUZ12 and the H3K9 methyltransferase SETDB1. The factors with the highest ROC AUC are CTCF and members of the cohesin complex, SMC3 and RAD21, that bind at CTCF sites.

**Figure 2:**
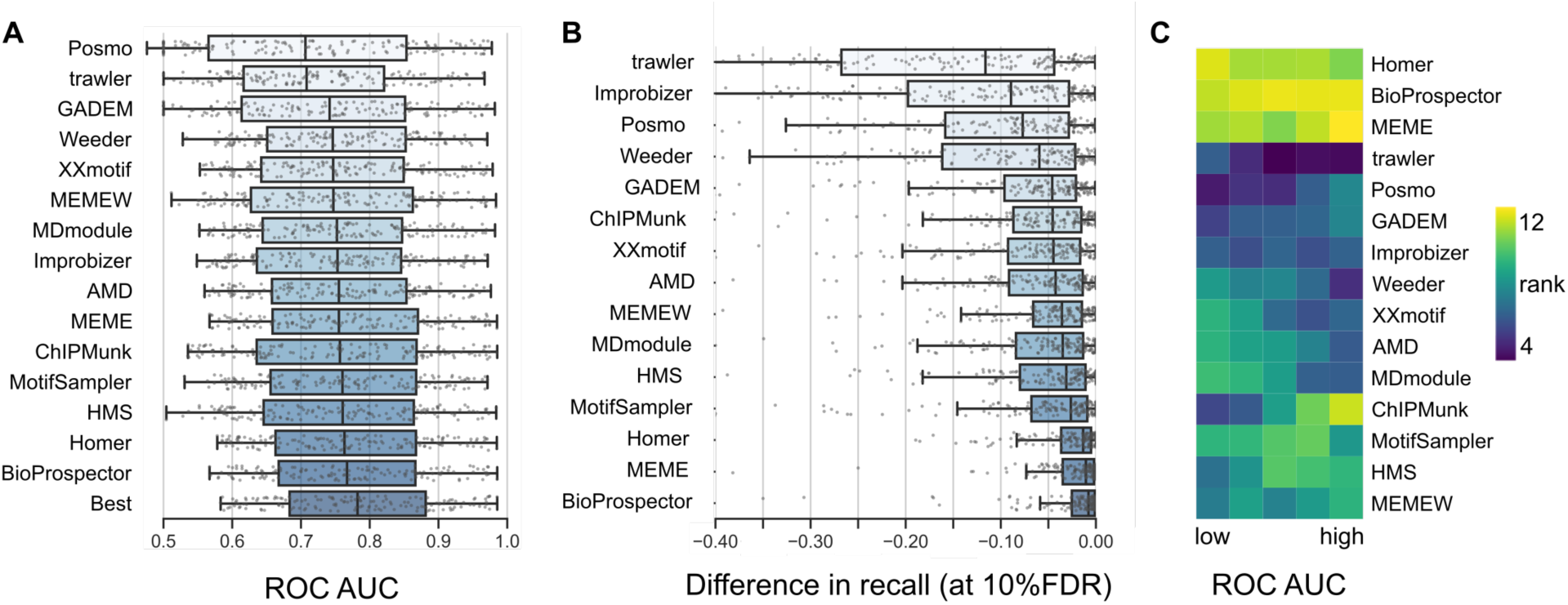
Benchmark of de novo motif finders. **A)** Comparison of the ROC AUC of the best motif of each motif finder. The boxplot shows the best motif per peak set of 495 peaks for 270 proteins from ENCODE. The best motif from all motif finders is indicated as ‘Best’. **B)** Comparison of the best motif per motif finder compared to the best overall motif for each data set. Plotted is the difference in recall compared to the best motif. Recall is calculated at 10% FDR. **C)** The relative motif rank as a function of the motif quality. Rank is the mean overall rank of three metrics (ROC AUC, recall at 10% FDR and MNCP).

Generally, the ROC AUC distribution of all evaluated motif finders is very similar. However, a few outliers can be observed. Trawler and Posmo show an overall lower distribution of ROC AUC scores. Compared to the ROC AUC scores of the next best program, GADEM, this is significant (p < 0.01, Wilcoxon signed-rank). Selecting the best motif for each experiment, as the ensemble method implemented in GimmeMotifs would do, results in a ROC AUC distribution that is significantly higher than the best single method, BioProspector (p < 1e-21, Wilcoxon signed-rank).

As stated in the previous section, the ROC AUC is not the best measure to evaluate motif quality. Therefore, we selected for every TF peak set the best overal motif on the basis of the recall at 10% FDR. We then plotted the difference between this best overall motif and the best motif from each individual *de novo* approach (Fig. 2B). For this figure, we used only the data sets where at least one motif had a recall higher than 0 at 10% FDR.

In line with previous results [18], there is no single tool that consistently predicts the best motif for each transcription factor. However, the motifs predicted by BioProspector, MEME and Homer are, on basis of this metric, consistently better than motifs predicted by other methods. In 75% of the cases, the motif predicted by BioProspector has a difference in recall smaller than 0.026 compared to the best overall motif. In this benchmark, four programs (Trawler, Improbizer, Posmo and Weeder) generally perform worse than average, with a mean decrease in recall of 0.11 to 0.17, as compared to the best motif. In addition, these programs tend to have a much more variable performance overall.

Predicted motifs identified using MEME with different motif widths show better performance than running MEME with the `minw` and `maxw` options (MEME vs. MEMEW in Fig. 2B). Of the best performing algorithms, both MEME and BioProspector were not specifically developed for ChIP-seq data, however, they consistently outperform most methods created for ChIP-seq data. Of the ChIP-seq motif finders Homer consistently shows good performance.

Finally, to gain further insight into *de novo* motif finder performance, we stratified the ChIP-seq datasets by motif “quality”. We divided the transcription factors into five bins on basis of the ROC AUC score of the best motif. For each bin we ranked the tools on basis of the average of three metrics (ROC AUC, recall at 10% FDR and MNCP [22]). The results are visualized as a heatmap in Figure 2C. From this visualization, it is again clear that BioProspector, MEME and Homer produce consistently high-ranking motifs, while the motifs identified by Trawler, Posmo, GADEM and Improbizer generally have the lowest rank. Interestingly, for some motif finders, there is a relation between motif presence and the relative rank. Weeder, XXMotif and MDmodule yield relatively high-ranking motifs when the ROC AUC of the best motif for the data set is low. On the other hand, ChIPMunk shows the opposite pattern. Apparently this algorithm works well when a motif is present in a significant fraction of the data set.

These results illustrate that motif finders need to be evaluated along a broad range of data sets with different motif presence and quality. Another interesting observation is that this ChIP-seq benchmark shows a lower-than-average performance for Weeder, which actually was one of the highest scoring in the Tompa et al. benchmark. It should be noted that our metric specifically evaluates how well *de novo* motif finders identify the primary motif in the context of high-scoring ChIP-seq peaks. It does not evaluate other aspects that might be important, such as the ability to identify low-abundant or co-factor motifs. Furthermore, with ChIP-seq data there are usually thousands of peaks available. This allows for other algorithms than those that work well on a few sequences. Interestingly, the original MEME shows consistently good performance, although the running time is longer than most other tools. On the basis of this analysis, BioProspector should be the top pick for a program to identify primary motifs in ChIP-seq data. However, an ensemble program such as GimmeMotifs will report high-quality motifs more consistently than any single tool.

### Differential motif analysis of hematopoietic enhancers identifies cell type-specific regulators

While many motif scanners and methods to calculate enrichment exist, there are few methods to compare motif enrichment or activity between two or more data sets. The CentriMo algorithm from the MEME suite implements a differential enrichment method to compare two samples [23]. Other approaches, such as MARA [24,25] and IMAGE [15], are based on linear regression. Here we present the *maelstrom* algorithm that integrates different methods to determine motif relevance or activity in an ensemble approach (Fig. 3A).

**Figure 3:**
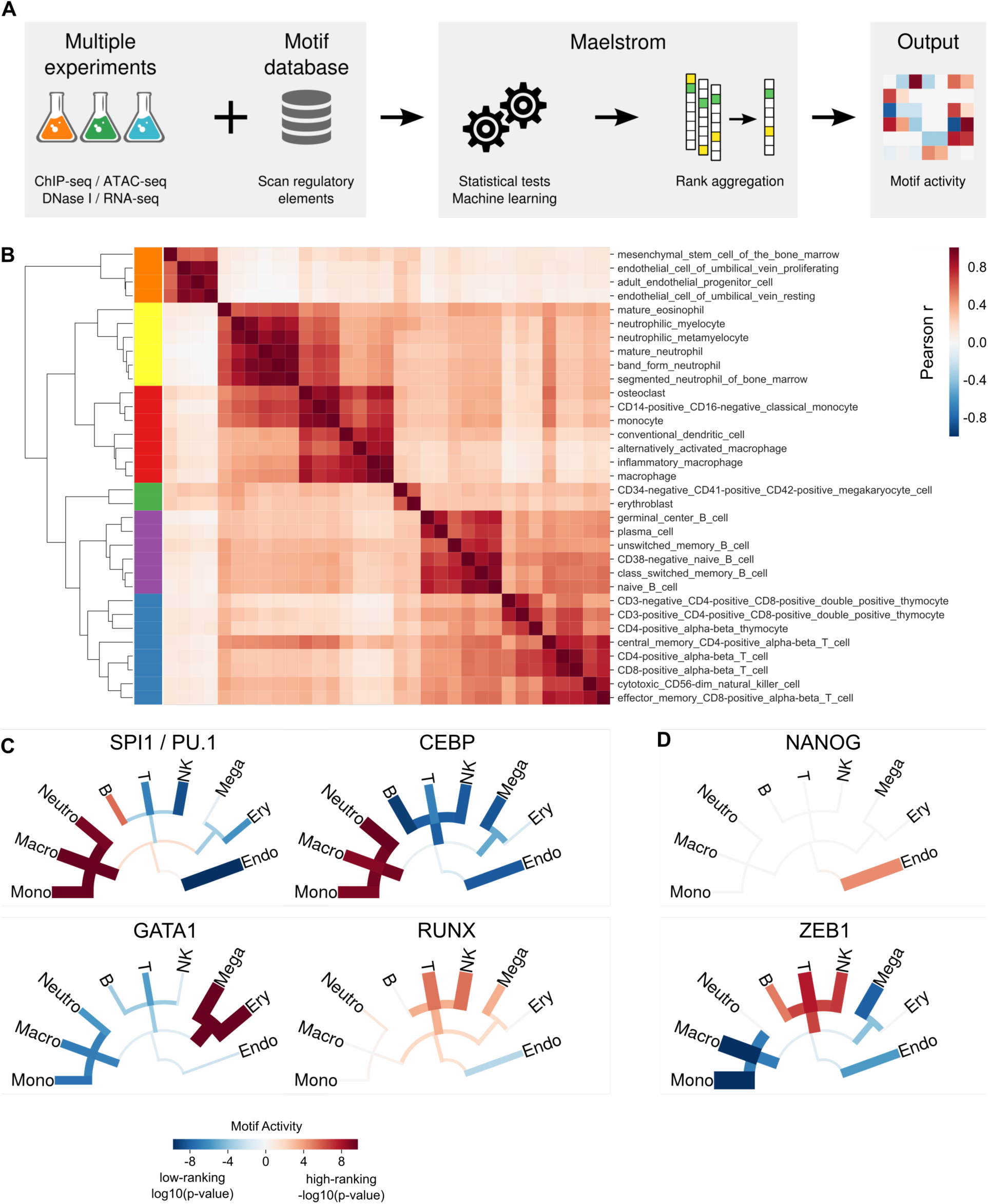
Predicting TF motif activity using maelstrom. **A)** A schematic overview of the maelstrom ensemble method. **B)** Heatmap of the correlation of H3K27ac signal in hematopoietic enhancers. We counted H3K27ac ChIP-seq reads in 2kb sequences centered at DNase I peaks. Counts were log2-transformed and scaled and replicates were combined by taking the mean value. This heatmap shows the Pearson r, calculated using the 50,000 most dynamic peaks. **C)** Selection of the results of running `gimme maelstrom` on the 50,000 most dynamic hematopoietic enhancers. Shown is the motif activity of four motifs associated with factors that are known to play a role in hematopoietic cells: SPI1 (PU.1), CEBP, RUNX and GATA1. The visualization shows a schematic, simplified cell lineage tree. The color and the line thickness represent the motif activity, where the value corresponds to the log10 of the p-value of the rank aggregation. For high-ranking motifs (red) -log10(p-value) is shown, while for low ranking motifs (blue) log10(p-value) of the reversed ranking is shown. **D)** Motif activity, as in **C**, of two motifs of factors for which the exact function in these cell types is currently unknown.

To demonstrate the utility of maelstrom we identified motif activity based on enhancers in hematopoietic cells. We downloaded 69 human hematopoietic DNaseI experiments (Supplementary Table S1), called peaks, and created a combined peak set as a collection of putative enhancers. In addition we downloaded 193 hematopoietic H3K27ac ChIP-seq experiments, mainly from BLUEPRINT [26] (Supplementary Table S1). We determined the number of H3K27ac reads per enhancer (Supplementary Table S2). After log2 transformation and scaling, we selected the 50,000 most dynamic peaks. Figure 3B shows the correlation of the H3K27ac enrichment in these 50,000 enhancers between cell types. For this plot, replicates were combined by taking the mean value and all experiments corresponding to treated cells were removed. We can observe six main clusters 1) non-hematopoietic cells 2) neutrophilic cells, 3) monocytes, macrophages and dendritic cells, 4) megakaryocytes and erythroblasts, 5) B cells 6) T cells and natural killer (NK) cells. We can conclude that the H3K27ac profile within this enhancer set recapitulates a cell type-specific regulatory signal.

To determine differential motif activity from these dynamic enhancers we used maelstrom. We combined Lasso, Bayesian ridge regression, multi-class regression using coordinate descent [27] and regression with boosted trees [28]. The coefficients or feature importances were ranked and combined using rank aggregation [29]. A p-value was calculated for consistently high ranking and consistently low ranking motifs. A selection of the results is visualized in Figure 3C. The full results are available as Supplementary File S1 and on Zenodo (10.5281/zenodo.1491482).

Two of the most signicant motifs are SPI1 (PU.1) and CEBP (Fig. 3C). The motif activity for SPI1 is high in monocytes and macrophages, consistent with its role in myeloid lineage commitment [30]. The CEBP family members are important for monocytes and granulocytic cells [31], and show a high motif activity in neutrophils and monocytes. Other strong motifs include RUNX for T cells and NK cells and GATA1 for erythroid cells (Fig. 3C).

We identified a high activity for motifs representing the ZEB1 and Snail transcription factors (Fig. 3D). The Snail transcription factors play an important role in the epithelial-to-mesenchymal transition (EMT), and their role in hematopoietic cells is less well-described. However, recently Snai2 and Snai3 were found to be required to generate mature T and B cells [32,33] in mice. ZEB1 is expressed in T cells and represses expression of IL-2 [34], as well as other immune genes such as CD4 [35] and GATA3 [36]. ZEB1 knockout mice exhibit a defect in thymocyte development [37]. Together, this suggests that these TFs could play an important role in the hematopoietic lineage.

Finally, an interesting observation is the predicted motif activity of NANOG in endothelial cells (Fig. 3D). NANOG is expressed in embryonic stem cells and is essential for maintenance of pluripotency [38]. However, NANOG is indeed also expressed in endothelial cells and has been shown to play a role in endothelial proliferation and angiogenesis [39].

In addition to the described examples, we identified a large compendium of TF motifs that display differential activity in the hematopoeitic lineage (Fig. 4). This demonstrates that `gimme maelstrom` can be used to analyze complex, multi-dimensional data sets such as this large-scale collection of hematopoietic enhancers. Especially in experiments where there are multiple conditions or time points that need to be compared, `maelstrom` provides a powerful method to determine differential transcription factor motif activity.

**Figure 4:**
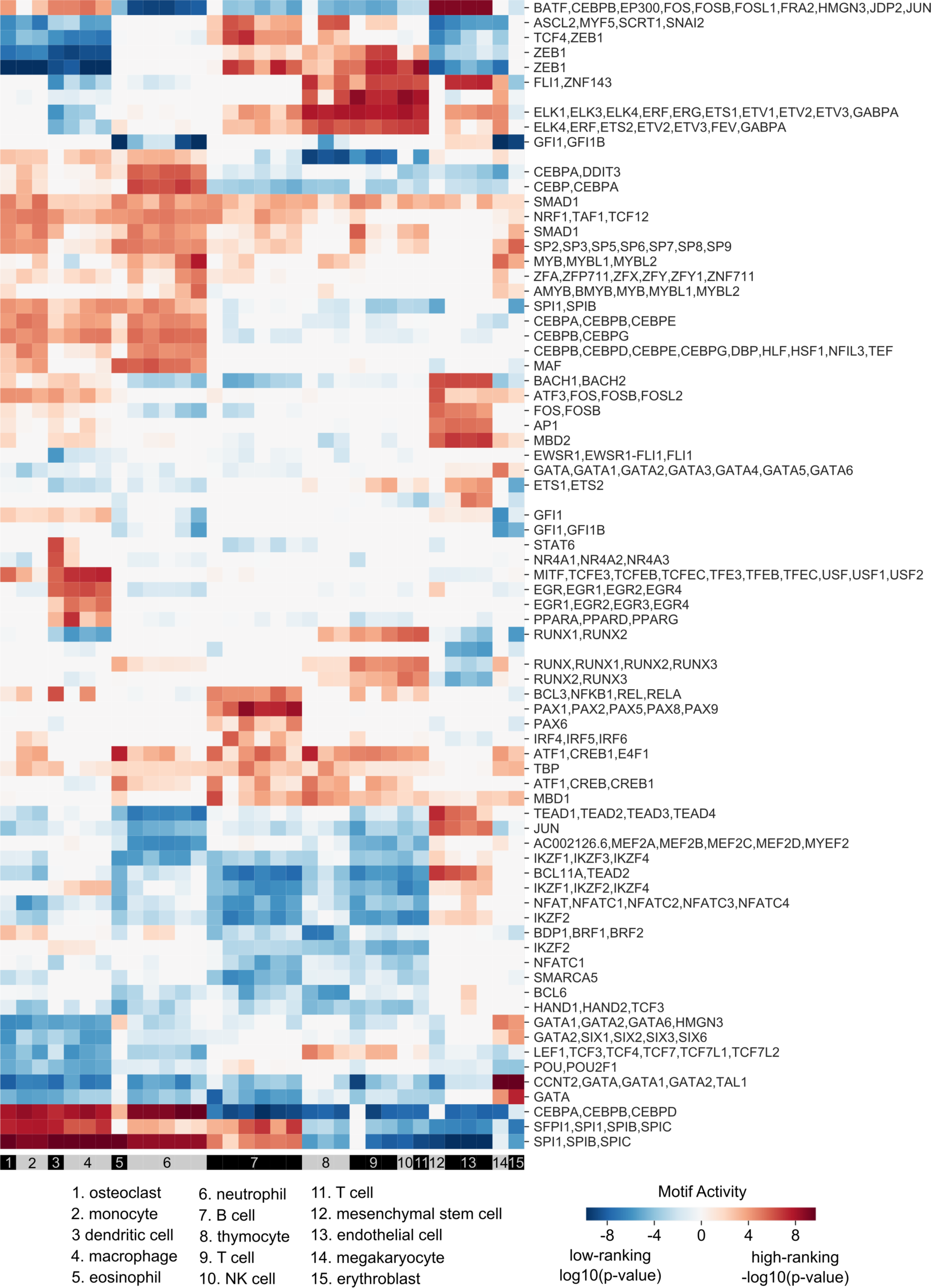
Predicting TF motif activity using maelstrom. Results of running `gimme maelstrom` on the 50,000 most dynamic hematopoietic enhancers. The motif activity of the top 78 motifs (absolute motif activity >= 5) is visualized in a heatmap. The color represents the reported motif activity, where the value corresponds to the log10 of the p-value of the rank aggregation. For high-ranking motifs (red) -log10(p-value) is shown, while for low ranking motifs (blue) log10(p-value) of the reversed ranking is shown.

## Methods

### GimmeMotifs

#### Implementation

GimmeMotifs is implemented in Python, with the motif scanning incorporated as a C module. The software is developed on GitHub (https://github.com/vanheeringen-lab/gimmemotifs/) and documentation is available at https://gimmemotifs.readthedocs.io. Functionality is covered by unit tests, which are run through continuous integration. GimmeMotifs can be installed via bioconda [40], see https://bioconda.github.io/ for details. All releases are also distributed through PyPi [41] and stably archived using Zenodo [42]. For *de novo* motif search, 14 different external tools are supported (Table 2). All of these are installed when conda is used for installation. By default, genomepy is used for genome management [43]. In addition, GimmeMotifs uses the following Python modules: numpy [44], scipy [45], scikit-learn, scikit-contrib-lightning [27], seaborn [46], pysam [47,48], xgboost [28] and pandas. In addition to the command line tools, all GimmeMotifs functionality is available through a Python API.

**Table 2:** External *de novo* motif prediction tools supported by GimmeMotifs. The version shown is the version string reported by the software. If the version is unknown, the date when the tool was added to the GimmeMotifs repository is shown.

| Name | Version | Citation |
| --- | --- | --- |
| AMD | Unknown (2012-11-21) | [49] |
| BioProspector | 4/15/04 | [50] |
| ChIPMunk | V7 10012017 | [51] |
| GADEM | v1.3.1 | [52] |
| HMS | Unknown (2012-11-23) | [53] |
| Homer | 2 | [6] |
| Improbizer | Unknown (2010-10-01) | [54] |
| MDmodule | Unknown (2010-07-29) | [55] |
| MEME | 4.6.0 | [56] |
| MotifSampler | 3.2 | [57] |
| Posmo | May-19-2011 | [58] |
| Trawler | 2.0 | [59] |
| Weeder | 2.0 | [60] |
| XXmotif | 1.6 | [61] |

#### De novo motif prediction pipeline

Originally, GimmeMotifs was developed to predict *de novo* motifs from ChIP-seq data using an ensemble of motif predictors [9]. The tools currently supported are listed in Table 2. An input file (BED, FASTA or narrowPeak format) is split into a prediction and validation set. The prediction set is used to predict motifs, and the validation set is used to filter for significant motifs. All significant motifs are clustered to provide a collection of non-redundant *de novo* motifs. Finally, significant clustered motifs are reported, along with several statistics to evaluate motif quality, calculated using the validation set. These evaluation metrics include ROC AUC, distribution of the motif location relative to the center of the input (i.e., the ChIP-seq peak summit) and the best match in a database of known motifs.

#### Motif activity by ensemble learning: maelstrom

GimmeMotifs implements eight different methods to determine differential enrichment of known motifs between two or more conditions. In addition, these methods can be combined in a single measure of *motif activity* using rank aggregation. Four methods work with discrete sets, such as different peak sets or clusters from a K-means clustering. The hypergeometric test uses motif counts with an empirical motif-specific FPR of 5%. All other implemented methods use the z-score normalized PFM log-odds score of the best match.

To combine different measures of motif significance or activity into a single score, ranks are assigned for each individual method and combined using rank aggregation based on order statistics [29]. This results in a probability of finding a motif at all observed positions. We use a Python implemention based on the method used in the R package RobustRankAggreg [62]. The rank aggregation is performed twice, once with the ranks reversed to generate both positively and negatively associated motifs.

The hypergeometric test is commonly used to calculate motif enrichment, for instance by Homer [6]. In GimmeMotifs, motifs in each cluster are tested against the union of all other clusters. The reported value is -log10(p-value) where the p-value is adjusted by the Benjamini-Hochberg procedure [63].

Using the non-parametric Mann-Whitney U test, GimmeMotifs tests the null hypothesis that the motif log-odds score distributions of two classes are equal. For each discrete class in the data, such as a cluster, it compares the score distributions of the class to the score distribution of all other classes. The value used as activity is the -log10 of the Benjamini-Hochberg adjusted p-value.

The two other methods are classification algorithms: random forest using scikit-learn and a large-scale multiclass classifier using block coordinate descent [27] as implemented in the scikit-contrib-lightning module. The classifier in GimmeMotifs uses a l1/l2 penalty with squared hinge loss where the alpha and C parameters are set using grid search in 10 fold cross-validation procedure.

The other four methods that are implemented relate motif score to an experimental measure such as ChIP-seq or ATAC-seq signal or expression level. These are all different forms of regression. In addition to ridge regression, which is similar to Motif Activity Response Analysis (MARA) [24,25], these methods include regression using boosted trees (XGBoost [28]), multiclass regression [27] and L1 regularized regression (LASSO).

### Clustering to create the gimme.vertebrate.v5.0 motif database

We collected all motifs from the following databases: CIS-BP v1.02 [12], ENCODE [4], Factorbook [3], HOCOMOCO v11 [5], HOMER v4.10 [6], JASPAR 2018 vertebrates [10] and SwissRegulon [13]. Motif similarity was calculated using Pearson correlation of motif scores profiles [64,65] using a sequence that contains each 7-mer or its reverse complement [66]. We then clustered the motifs using agglomerative clustering with complete linkage and connectivity constraints where only motifs with a Pearson r >= 0.5 were considered as neighbors. The number of clusters was set to 1900. After clustering, we discarded all motifs were the sum of the information content of all positions was less than 5. The clustered database, `gimme.vertebrate.v5.0`, is distributed with GimmeMotifs.

### Transcription factor motif database benchmark

We downloaded all hg38 non-ENCODE ChIP-seq peaks from Remap 2018 v1.2 [16] (http://tagc.univ-mrs.fr/remap/index.php?page=download). We removed all factors with fewer than 1000 peaks and created regions of 100 bp centered at the peak summit. Background files were created for each peak set using bedtools shuffle [67], excluding the hg38 gaps and the peak regions. The ROC AUC and Recall at 10% FDR statistics were calculated using `gimme roc`. The motif databases included in the comparison are listed in Table 3. We only included public databases that can be freely accessed and downloaded.

**Table 3:** Motif databases.

| Name | Version | Citation |
| --- | --- | --- |
| Factorbook | Sep. 2012 | [3] |
| SwissRegulon | Nov. 2006 | [13] |
| GimmeMotifs vertebrate | v5.0 |  |
| HOCOMOCO | v11 | [5] |
| Homer | v4.10 | [6] |
| JASPAR | 2018 | [10] |
| ENCODE | Dec. 2013 | [4] |
| Madsen | 1.1 | [15] |
| RSAT vertebrate clusters | Sep. 2017 | [14] |

### De novo motif prediction benchmark

We downloaded all spp ENCODE peaks (January 2011 data freeze) from the EBI FTP (http://ftp.ebi.ac.uk/pub/databases/ensembl/encode/integration_data_jan2011/byDataType/peaks/jan2011/spp/optimal/). We selected the top 5000 peaks and created 100bp regions centered on the peak summit. As background we selected 100 bp regions flanking the original peaks. For the de novo motif search default settings for `gimme motifs` were used. The workflow is implemented in snakemake [68] and is available at https://github.com/vanheeringen-lab/gimme-analysis.

### Motif analysis of hematopoietic enhancers

To illustrate the functionality of `gimme maelstrom` we analyzed an integrated collection of hematopoietic enhancers. We downloaded all H3K27ac ChIP-seq and DNase I data from BLUEPRINT and hematopoietic DNase I data from ROADMAP (Supplementary Table S1). All DNase I data were processed using the Kundaje lab DNase pipeline version 0.3.0 https://github.com/kundajelab/atac_dnase_pipelines [69]. The ChIP-seq samples were processed using the Kundaje lab AQUAS TF and histone ChIP-seq pipeline https://github.com/kundajelab/chipseq_pipeline. For all experiments from BLUEPRINT we used the aligned reads provided by EBI. All ROADMAP samples were aligned using bowtie2 [70] to the hg38 genome. DNase I peaks were called using MACS2 [71] according to the default settings of the DNase pipeline. We merged all DNase I peak files and centered each merged peak on the summit of the strongest individual peak. H3K27ac reads were counted in a region of 2kb centered at the summit (Supplementary Table S2) and read counts were log2-transformed and scaled. We removed all samples that were treated and averaged all samples from the same cell type. We then selected all enhancers with at least one sample with a scaled log2 read count of 2, sorted by the maximum difference in normalized signal between samples and selected the 50,000 enhancers with the largest difference. Using this enhancer collection as input, we ran `gimme maelstrom` using default settings. The motif analysis workflow is implemented in a Jupyter notebook and is available at https://github.com/vanheeringen-lab/gimme-analysis.

## Conclusions

We demonstrated the functionality of GimmeMotifs with three examples. First, to evaluate different public motif databases, we quantified their performance on distinguishing ChIP-seq peaks from background sequences. The databases that perform best on this benchmark are collections of motifs from different sources. Of the individual databases HOCOMOCO and Factorbook rank highest using this collection of human ChIP-seq peaks. Based on our results it is recommended to use a composite database, such as the RSAT clustered motifs or the GimmeMotifs database (v5.0), for the best vertebrate motif coverage. However, these motifs are less well annotated. For instance, motifs based on ChIP-seq peaks from some sources might be from co-factors or cell type-specific regulators instead of the factor that was assayed. An example are motifs that are associated with the histone acetyl tranferase EP300. This transcriptional co-activator lacks a DNA binding domain, and associated motifs depend on the cell type. For instance, in a lymphoblastoid cell line such as GM12878 these include PU.1 and AP1. The lack of high-quality annotation makes it more difficult to reliably link motifs to transcription factors. This can be a distinct advantage of a database such as JASPAR. Although the motifs might not be optimal for every TF, JASPAR contains high-quality metadata that is manually curated.

In the second example, we benchmarked 14 different *de novo* motif finders using a large compendium of ChIP-seq data. While performance can vary between different data sets, there are several *de novo* motif finders that consistently perform well, with BioProspector, MEME and Homer as top performers. Interestingly, only Homer was specifically developed for ChIP-seq data. An ensemble approach such as GimmeMotifs still improves on the use of individual tools. This example also illustrates that newly developed *de novo* motif finders should be evaluated on many different data sets, as this is necessary to accurately judge the performance.

Finally, we presented a new ensemble approach, *maelstrom*, to determine motif activity in two or more epigenomic or transcriptomic data sets. Using H3K27ac ChIP-seq signal as a measure for enhancer activity, we analyzed cell-type specific motif activity in a large collection of hematopoietic cell types. We identified known lineage regulators, as well as motifs for factors that are less well studied in a hematopoietic context. This illustrates how `gimme maelstrom` can serve to identify cell type-specific transcription factors and has the potential to discobver

In conclusion, GimmeMotifs is a flexible and highly versatile framework for transcription factor motif analysis. Both command line and programmatic use in Python are supported. One planned future improvement to GimmeMotifs is the support of more sophisticated motif models. Even though the PFM is a convenient representation, it has certain limitations. A PFM cannot model inter-nucleotide dependencies, which are known to affect binding of certain TFs. Multiple different representations have been proposed [72,73,74,75,76], but no single one of these has gained much traction. It is still unclear how well these models perform and their use depends on specific tools. Supporting these different models and benchmarking their performance relative to high-quality PFMs will simplify their use and give insight into their benefits and disadvantages. Second, there is significant progress recently in modeling TF binding using deep neural networks (DNNs) [77,78]. These DNNs can learn sequence motifs, as well as complex inter-dependencies, directly from the data. However, while biological interpretation is possible [79], it becomes less straightforward. We expect that analyzing and understanding a trained DNN can benefit from high-quality motif databases and comparative tools such as GimmeMotifs.

## Availability and requirements

- Project name: GimmeMotifs
- Project home page: https://github.com/vanheeringen-lab/gimmemotifs
- Operating system(s): Linux, Mac OSX
- Programming language: Python 3
- Other requirements: *de novo* motif finders
- License: MIT

## Availability of supporting data

- Scripts and notebooks to reproduce the analysis are available at:
  - https://github.com/vanheeringen-lab/gimme-analyis.
- Additonal data files are available at Zenodo:
  - Table of H3K27ac read counts at all DNase I accessible sites: 10.5281/zenodo.1488669.
  - Results of gimme maelstrom (Figure 3 and 4): 10.5281/zenodo.1491482.

## Additional files

- Supplementary Table S1: List of accessions used in the analysis of hematopoietic enhancers.
- Supplementary Table S2: Table of H3K27ac read counts at all DNase I accessible sites.
- Supplementary File S1: Results of gimme maelstrom (Figure 3 and 4).

## Author contributions

SJvH designed and performed experiments and analysis, wrote the code and wrote the manuscript. NB processed and analyzed the hematopoietic DNase I and H3K27ac ChIP-seq data. All authors read and approved the final manuscript.

## Acknowledgements

We would like to thank all research symbionts, who make their tools and data publicly available. This study makes use of data generated by the Blueprint Consortium. A full list of the investigators who contributed to the generation of the data is available from www.blueprint-epigenome.eu. Additionally, this study used data provided by the NIH Roadmap Epigenomics Consortium (http://nihroadmap.nih.gov/epigenomics/).

